# Negative interplay between biofilm formation and competence in the environmental strains of *Bacillus subtilis*

**DOI:** 10.1101/2020.06.15.153833

**Authors:** Qianxuan She, Evan Hunter, Yuxuan Qin, Samantha Nicolau, Eliza A. Zalis, Hongkai Wang, Yun Chen, Yunrong Chai

## Abstract

Environmental strains of the soil bacterium *Bacillus subtilis* have valuable applications in agriculture, industry, and biotechnology. They are capable of forming robust biofilms and demonstrate excellent biological control activities in plant protection. However, environmental strains are genetically less accessible, a sharp contrast to the laboratory strains well known for their natural competence and a limitation toward their application. In this study, we observed that robust biofilm formation of the environmental strains greatly reduces the rate of competent cells within the biofilm. By using the model strain 3610, we reveal a cross-pathway regulation that allows biofilm matrix producers and competence-developing cells to undergo mutually exclusive cell differentiation. We show that the competence activator ComK represses the key biofilm regulatory gene *sinI* by directly binding to the *sinI* promoter, thus blocking competent cells from simultaneously becoming matrix producers. In parallel, the biofilm activator SlrR represses competence through three distinct mechanisms, involving both genetic regulation and cell morphological changes. We discuss potential implications of limiting competence in a bacterial biofilm.

**Importance:** The soil bacterium *Bacillus subtilis* is capable of forming robust biofilms, a multicellular community important for its survival in the environment. *B. subtilis* also exhibits natural competence, the ability of cells to acquire genetic materials directly from the environment. By investigating competence development *in situ* during *B. subtilis* biofilm formation, we reveal that robust biofilm formation, an important feature of the environmental strains of *B. subtilis*, often greatly reduces the rate of competent cells within the biofilm. We characterize a cross-pathway regulation that allows cells associated with these two developmental events to undergo mutually exclusive cell differentiation during biofilm formation. Finally, we discuss potential biological implications of limiting competence in a bacterial biofilm.

## Introduction

*Bacillus subtilis* is a soil-dwelling, spore-forming bacterium widely present in nature, and plays important roles in environment, agriculture, and industry. *B. subtilis* is also a plant growth-promoting rhizobacterium (PGPR), and a biological control agent that shows various beneficial activities in plant protection (1). Biological control by *B. subtilis* is attributed to a number of important abilities of the bacterium, including antibiotic production, inhibition of pathogenic fungi and parasites, induction of plant systemic resistance, and formation of plant root-associated biofilms (2-4). Biofilms are communities of microorganisms encased in a self-produced extracellular matrix, which provides protection to cells within the biofilm against various biotic and abiotic stresses (5-8). Undomesticated environmental strains of *B. subtilis* form robust biofilms in response to a variety of environmental and cellular signals (9-16). Some of those strains, such as NCIB3610 (hereafter abbreviated as 3610), become a model system to study bacterial biofilm formation (2, 14, 17-20).

In *B. subtilis*, biofilm formation is initiated by one or several histidine kinases (KinA-KinD) sensing environmental signals and activating a phosphorelay (Spo0F-Spo0B-Spo0A), which then activates Spo0A, a master regulator for sporulation and biofilm formation, via protein phosphorylation (Fig. 1)(21, 22). Phosphorylated Spo0A (Spo0A~P) activates *sinI*; the product SinI is an antagonist that counteracts a biofilm repressor SinR (23, 24). Another regulatory protein SlrR is also involved in antagonizing SinR (25). SlrR and SinR form a double negative regulatory switch (26). Under non-biofilm conditions, those matrix operons are repressed. Interestingly, domesticated strains of *B. subtilis* that have been applied in the laboratory research for multiple decades due to their feasibility in genetic manipulation, have lost their ability to form robust biofilms due to genetic mutations occurred during domestication (19, 20, 27).

**Figure 1.**
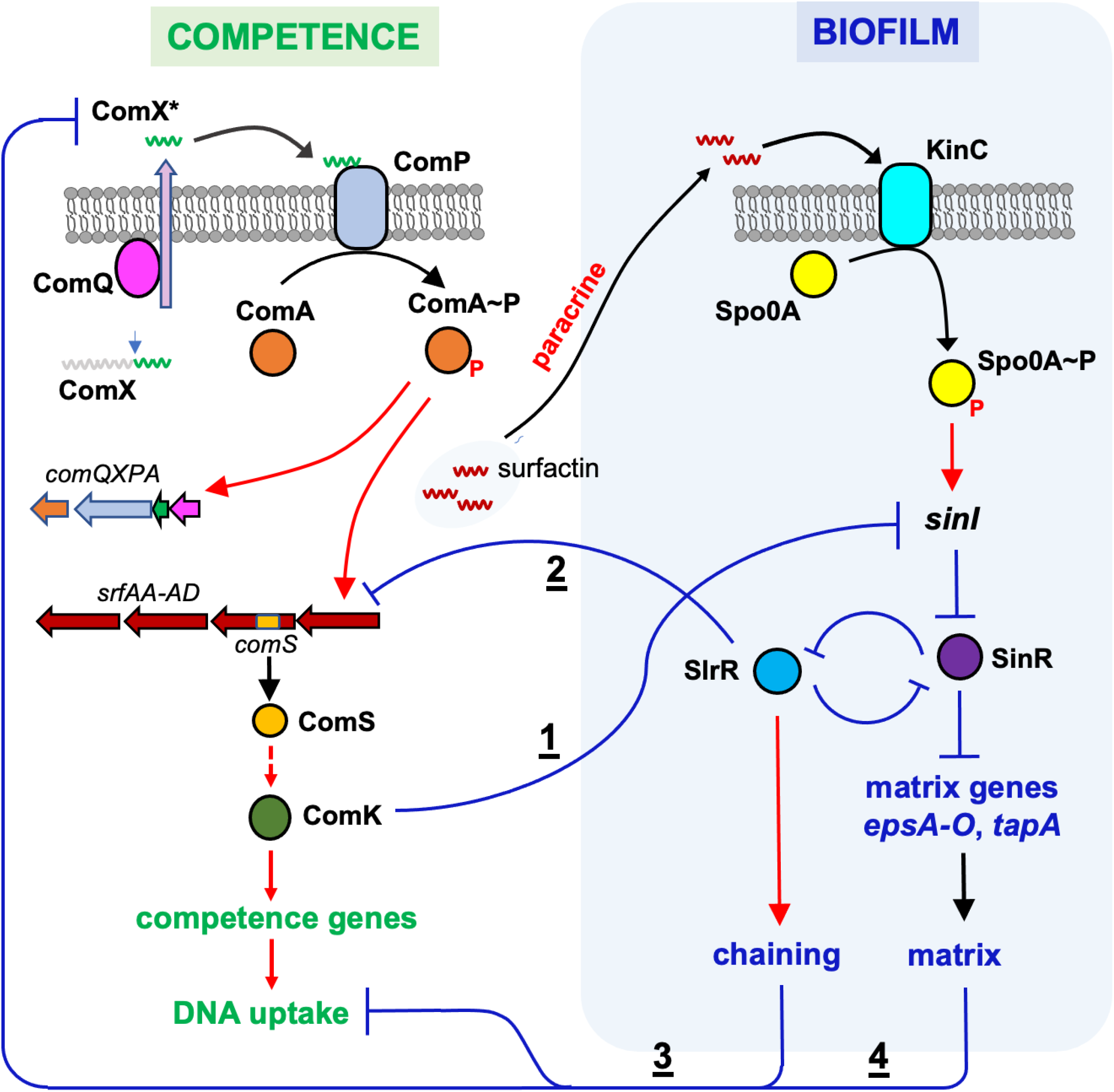
A working model for cross-pathway regulation between competence and biofilm in *B. subtilis*. Competence development initiates when the quorum-sensing (QS) peptide derived from ComX (ComX*) is sensed by a membrane histidine kinase ComP (37). The response regulator ComA then activates a *srfAA-AD* operon and an embedded small gene *comS;* the latter encodes a positive regulator ComS for the competence activator ComK (42). ComK^ON^ cells express late competence genes, which ultimately differentiate into competent cells ready for DNA uptake. Here we propose that ComK simultaneously and negatively regulates the biofilm pathway by repressing the key biofilm regulatory gene *sinI* (shown as 1). SinI antagonizes the biofilm master repressor SinR to derepress genes for the biofilm matrix production (*epsA-O, tapA, etc*). Negative regulation of *sinI* by ComK is expected to inhibit biofilm formation. SlrR is another antagonist of SinR and forms a double negative loop with SinR. Under biofilm inducing conditions, *sinI* is activated by the developmental master regulator Spo0A in response to sensory kinases (e.g. KinC) sensing various environmental signals. Here, we also propose that the biofilm regulator SlrR negatively regulates competence development through several distinct mechanisms. First, SlrR activates matrix production, which physically blocks sensing of the quorum-sensing peptide signal ComX* (shown as <u>4</u>)(49); second, SlrR-induced cell chaining may block DNA uptake since DNA uptake machinery was shown to be pole-localized (57, 58)(shown as <u>3</u>); third, SlrR negatively regulates the *srfAA-AD* operon and *comS* (shown as <u>2</u>). Red arrows and blue lines represent positive and negative regulation, respectively. ComX*, a secreted QS peptide derived from ComX. Surfactin induces matrix production by a paracrine signaling mechanism (49).

Cell differentiation is another hallmark feature in bacterial biofilm formation. In *B. subtilis*, it is known that cells in the biofilm differentiate into phenotypically distinct cell types (2, 14, 28). Some of these cell types may overlap or become mutually exclusive. Previous studies showed that a subpopulation of cells progressed to become sessile matrix producers while cells in another subpopulation remained motile (26, 29). The two subsets of cells are mutually exclusive due to the control via an epigenetic switch involving several regulatory genes including *sinI, sinR, slrR*, and *ymdB* (26, 30-32). The switch allows bifurcation of the population into SlrR^ON^ and SlrR^OFF^ cells, which correspond to sessile and motile cells due to SlrR-mediated gene regulation (26, 32). Some cells in the *B. subtilis* biofilm undergo sporulation (19). It was found that heat-resistant spores constitute 15-20% of the total cell population in a mature *B. subtilis* biofilm. It was proposed that in the matrix producers, Spo0A~P accumulates to intermediate levels to induce biofilm development (33). When Spo0A~P activities keeps rising, it eventually activates expression of hundreds of genes involved in sporulation (34, 35). This may explain how matrix producers transition to become sporulating cells and why sporulating cells, as a cell type, are inclusive to matrix producers. Competent cells are those capable of acquiring DNAs from the environment and are believed to be present in the *B. subtilis* biofilm as well (28). However, competence development has not been well studied *in situ* in a *B. subtilis* biofilm (36).

Natural competence is an ability of certain bacterial species, such as *B. subtilis*, to acquire environmental DNAs for genetic exchange (37, 38). Natural competence is evolutionarily important for the bacteria to increase genetic diversity and adaptability. In *B. subtilis*, the complex regulatory network controlling competence development has been elegantly elucidated (37-40). Competence is initiated when cells produce a quorum-sensing molecule, a peptide pheromone derived from ComX (Fig. 1)(41). This quorum-sensing peptide is sensed on the membrane by a sensory kinase ComP of the ComA-ComP two-component system (37). The response regulator ComA then activates a *srfAA-AD* operon, which not only encodes enzymes involved in biosynthesis of surfactin, but surprisingly also transcribes a small gene named *comS* (42). The protein ComS is critical in activating the competence activator ComK by helping release of ComK from a protein called MecA, an adaptor protein that normally routes ComK to be degraded (43, 44). Hence, the ComS-mediated release of ComK from MecA allows ComK to accumulate. In addition, the *comK* gene is subject to a bistable control mechanism; only a subset of cells accumulate high levels of this competence activator and enter the so-called K-state (39). Different from the well-studied laboratory strains, many environmental strains of *B. subtilis* are much less competent in general for still unclear reasons. This may limit the application of those environmental strains in industry and agriculture. In the model strain 3610, a *comI* gene located in a cryptic plasmid was shown to inhibit the competence; deletion of *comI* or cure of the cryptic plasmid boosted transformation efficiency of 3610 by more than one hundred fold (45). Whether such a plasmid-born competence inhibitory gene is broadly present in environmental strains is not known. Another regulatory gene *degQ* was also shown to reduce competence in 3610; deletion of *degQ* similarly boosted transformation efficiency of 3610 (27). In addition, there was convincing evidence that a point mutation in the promoter of *degQ* in the laboratory strain 168, which likely lowers the expression of *degQ*, contributes to the much higher transformation efficiency of 168 (27).

In this study, we aim to investigate factors that impact competence in the environmental strains of *B. subtilis*. We reveal an interplay between biofilm formation and competence development; robust biofilm formation in the environmental strains greatly reduces the rate of competent cells within the biofilm. We show that a very low number of cells express a late competence gene reporter and that those cells become mutually exclusive from matrix producers in the biofilm. We characterize a cross-pathway regulation that contributes to the above mutual exclusivity and limits competence in individual cells in the *B. subtilis* biofilm.

## Results

### *B. subtilis* environmental strains are robust biofilm formers but poor in competence

We previously investigated a number of environmental isolates of *B. subtilis* for their biological control activities in plant protection (4). Many of those strains form robust biofilms both under the laboratory setting (Fig. 2A) and on plant roots (4, 10). On the other hand, most of those environmental strains are much more difficult to manipulate genetically than the laboratory strains. They showed a much lower transformation efficiency, hundreds to tens of thousands fold lower than that of the laboratory strain 168 (Fig. 2B); some were not even transformable (data not shown). In those tested strains, big variations in transformation efficiency were also seen. For example, there was a several hundred-fold difference in transformation efficiency between 3610 and CY54, as judged by the percentage of transformants relative to the total number of cells (Fig. 2B). To test if variations in transformation efficiency were due to altered competence gene regulation, we constructed a fluorescent reporter for a late-stage competence gene *comGA* (P*_comGA_-gfp*), and introduced the reporter into those environmental strains as well as the laboratory strain 168. The engineered reporter strains were grown in the competence medium (MC) to early stationary phase, and cells were examined under fluorescent microscopy. P*_comGA_-gfp* expressing cells were observed in only a small subset of cells in the population (with the exception of 168, Fig. 3A). A clear bimodal pattern in P*_comGA_-gfp* expression was also seen, indicating that bistability in competence development reported previously in laboratory strains is also reinforced in environmental strains (39). The ratio of P*_comGA_-gfp* expressing cells in different environmental strains varied significantly and ranged from about 0.25% to 7.7%, much lower than in 168, the ratio in which was seen ~40% in our hand (Fig. 3B). In general, the results from assays of the fluorescent reporter correlated well with those from genetic transformation with the exception of CY54 (Figs. 2B and 3B), suggesting that the reduced transformation efficiency in environmental strains is likely due to genetic regulation. In summary, those environmental strains are robust biofilm formers but poor in competence.

**Figure 2.**
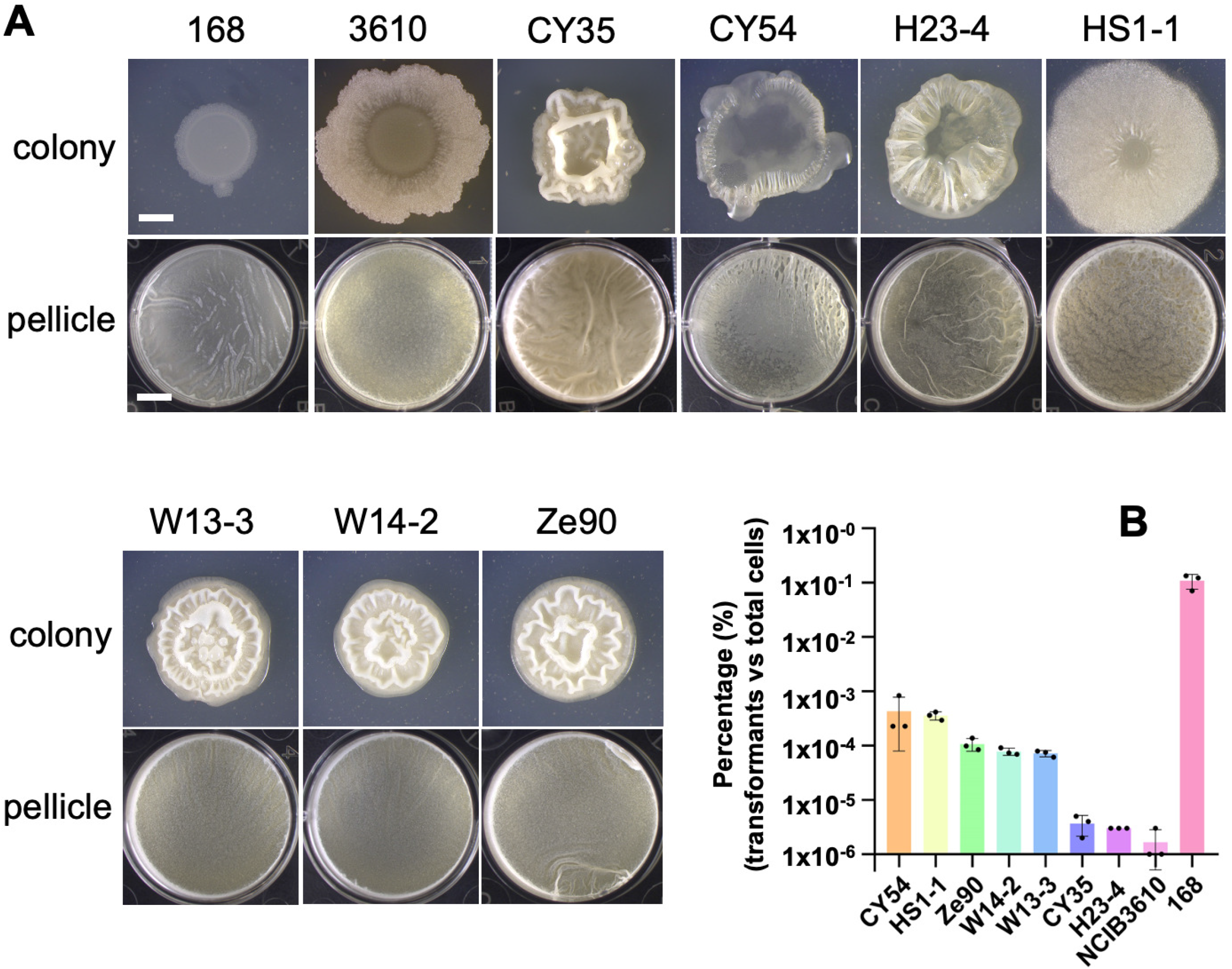
*B. subtilis* environmental strains are strong biofilm producers but poor in competence. **(A)** The colony and pellicle biofilm phenotypes of 7 environmental strains of *B. subtilis* plus 168 and 3610. Scale bar in the picture of colony, 2 mm; scale bar in the picture of pellicle, 5 mm. The scale bar in the picture of colony is representative for all pictures of colonies, so as the scale bar in the picture of pellicle. **(B)** Transformation efficiency of the 7 environment isolates of *B. subtilis* plus 168 and 3610. Results are shown as percentage (%) of the number of transformants relative to the total number of cells. Assays were performed in triplicates. Each dot represents one technical replicate. Error bars represent standard deviations.

**Figure 3.**
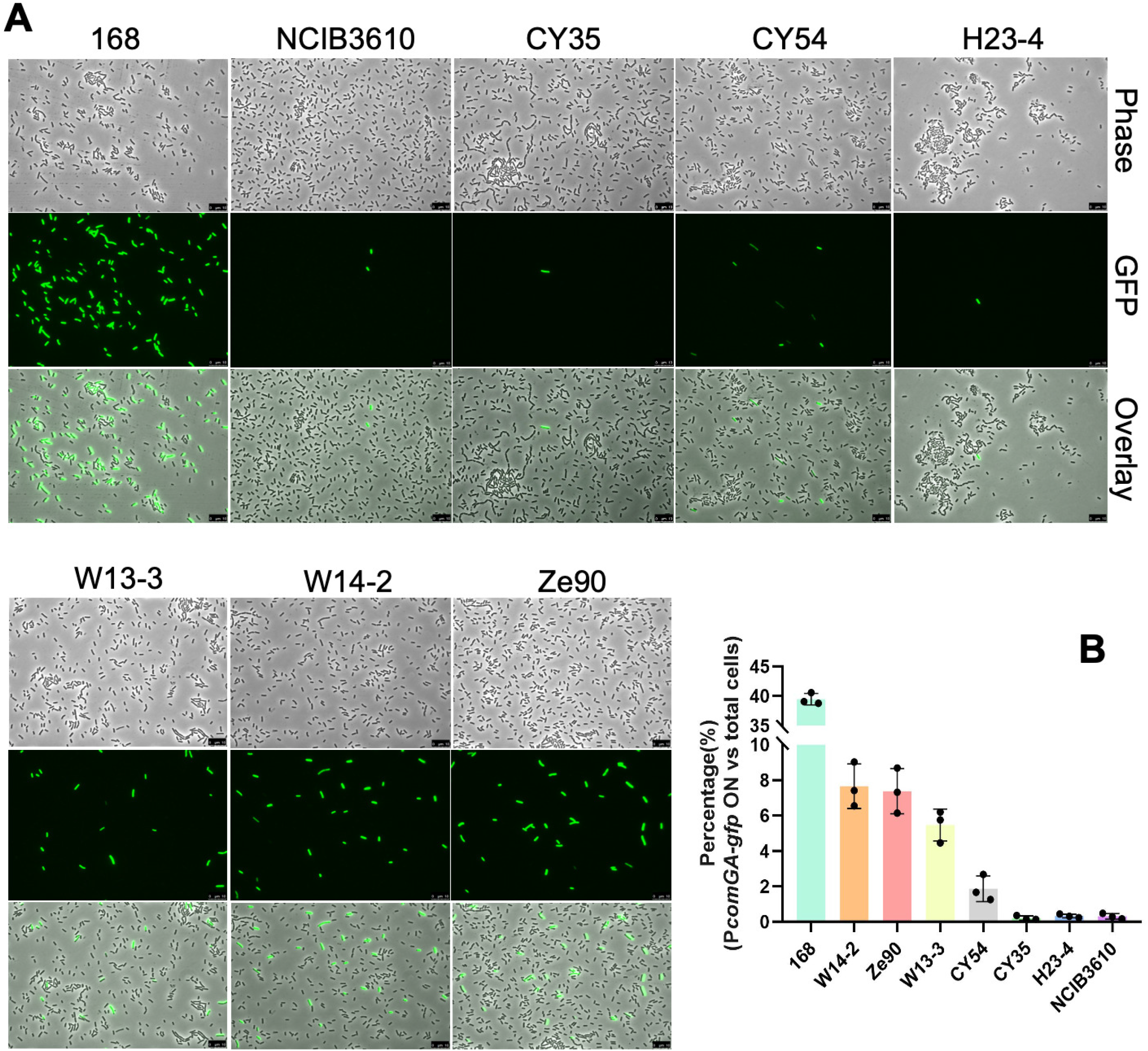
A small proportion of cells from the environmental strains express the late competence gene *comGA*. **(A)** Environmental strains harboring the late competence gene reporter P*_comGA_-gfp* were grown in the competence medium (MC) to early stationary phase. Cells were harvested and observed under fluorescent microscopy. The 168 and 3610 strains were included for comparison. Representative images were shown here. Scale bars, 10 μm. **(B)** The percentage of P*_comGA_-gfp* expressing cells relative to the total number of cells in 7 different environmental strains plus 168 and 3610. In each bar, three dots represent three individual data points calculated from three different images comprising of about 600-800 cells in total per sample. Error bars represent standard deviations.

### DegQ negatively impacts competence in some, but not all, tested environmental strains

Previous studies suggested that the *degQ* gene negatively regulates genetic competence in *B. subtilis* (27). A deletion mutation in *degQ* increased the transformation efficiency of 3610 by a few fold while *degQ* overexpression led to a reduction in the transformation efficiency (Supple. Fig. 1A). DegQ is believed to impact competence through DegU, a response regulator and a transcription factor on the *comK* gene. A *degU* deletion mutation almost completely eliminated competence in 3610, while the deletion mutation of *degS*, which encodes the histidine kinase of the DegS-DegU two-component system (46), modestly impaired the transformation efficiency (Supple. Fig. 1A).

To test if DegQ plays a similar role in competence in the environmental strains, we introduced Δ*degQ* into those strains and examined the transformation efficiency of the resulting mutants. Δ*degQ* increased transformation efficiency in 5 out of the 7 tested environmental strains (except for Ze90 and W13-3). The increase ranged from ~2 to 88 fold when compared to the wild type strains (Supple. Fig. 1B). In Ze90 and W13-3, Δ*degQ* decreased the transformation efficiency by ~4 and 7 fold, respectively (Supple. Fig. 1B). This result indicates that the impact of DegQ on competence varies in different environmental strains. We also wondered if the same or similar point mutation identified in the *degQ* promoter in the laboratory strain 168 is present in any of those environmental strains. We amplified the promoter region of the *degQ* gene by PCR and performed DNA sequencing. Our sequencing results show that the point mutation in *degQ* in 168 is not present in any of the environmental strains (highlighted in the blue square, Supple. Fig. 1C). However, additional mutations were identified in the promoter of *degQ* in CY35 and CY54 (single nucleotide changes at −28, −48, and −77 positions from the *degQ* transcription start, highlighted in red squares, Supple. Fig. 1C). Whether these newly identified mutations impact *degQ* expression and influence competence in CY54 and CY35 is yet to be tested.

### Matrix producers and competent cells are mutually exclusive in the *B. subtilis* biofilm

To investigate why many environmental strains are poor in competence, we decided to examine competence development *in situ* during biofilm formation in the model strain 3610. A dual-labelled fluorescent reporter strain of 3610 (P_*tapA*_-*mkate2* and P*_comGA_-gfp*, EH43) was constructed, which allowed us to measure the expression of the matrix operon *tasA-sipW-tapA* (P_*tapA*_-*mKate2*) and the activity of the late stage competence gene *comGA* (P*_comGA_-gfp*) simultaneously in the same cells (47, 48). Cells from a 3-day pellicle biofilm by the reporter strain were collected and examined under fluorescent microscopy. Cells were seen in bundled chains and showed strong expression of the matrix reporter P_*tapA*_-*mKate2* [chaining and matrix production are known to be coregulated during biofilm development, (32)](Fig. 4A and Supple. Fig. 2). In contrast, cells expressing P*_comGA_-gfp* were very rare, always in singlets, and almost never overlapped with cells expressing P_*tapA*_-*mKate2*. Since the number of cells expressing P*_comGA_-gfp* was very low, flow cytometry was applied to quantitatively determine the ratio of cells expressing the two reporters in the pellicle biofilm. Cells were similarly collected, treated with mild sonication to disrupt the bundled chains, and applied to flow cytometry. As shown in Fig. 4B-C, only 0.08% of the total cells expressed P*_comGA_-gfp* (Average results from three replicates were shown in Fig. 4C). The reason why the ratio of P*_comGA_-gfp* expressing cells was even lower in this assay than previously observed in Fig. 3B (~0.25% for 3610) is likely because the assay in Fig. 3 was done under competence favorite conditions (use of competence medium, cells collected at the early stationary phase, *etc*). More importantly, the results shown here (Fig. 4A-B and Supple. Fig. 2) strongly suggest that in the *B. subtilis* biofilm, matrix producers and competent cells rarely overlap; they are mutually exclusive cell types.

**Figure 4.**
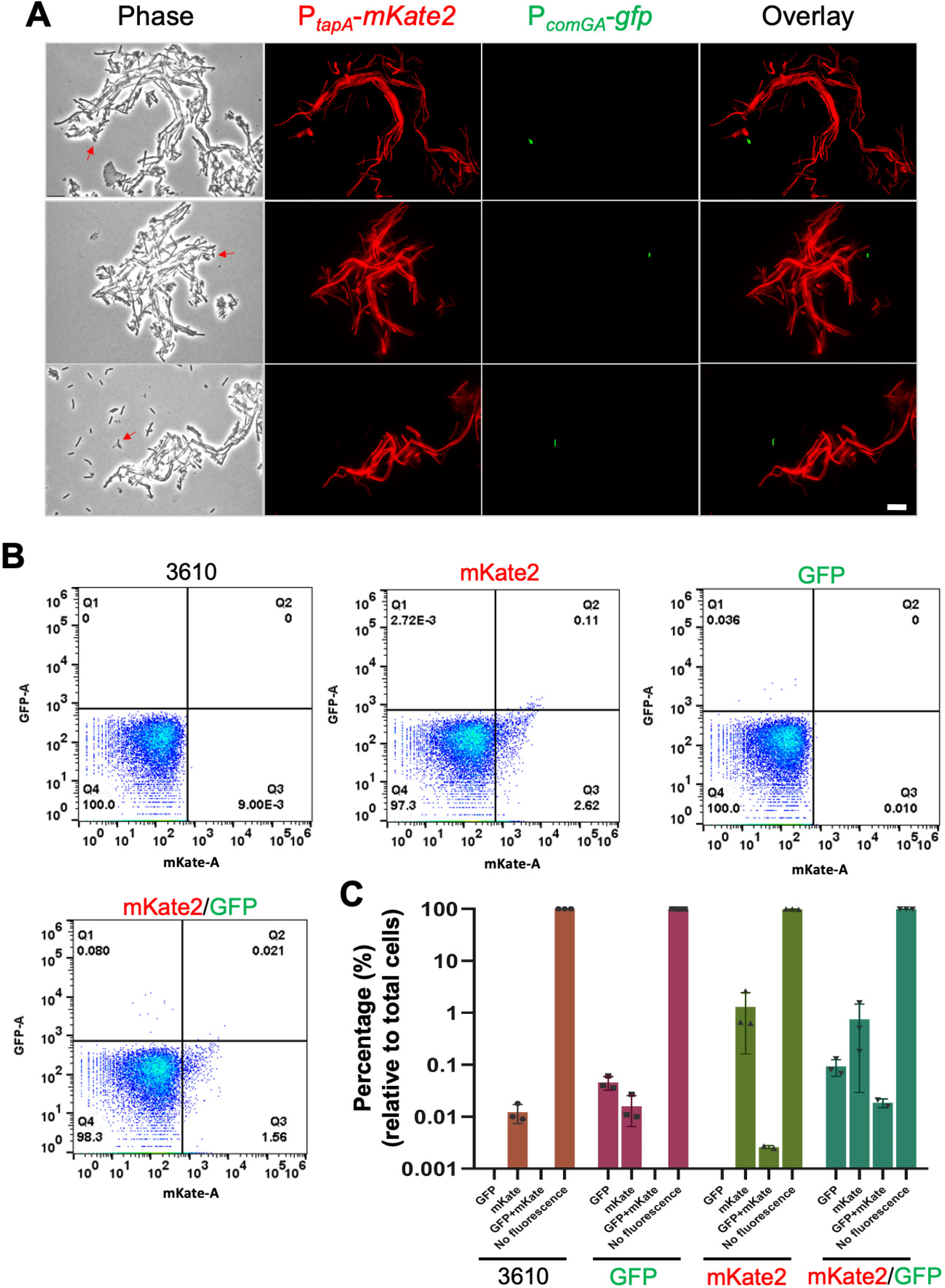
Matrix producers and competent cells are mutually exclusive in the 3610 biofilm. **(A)** Fluorescent microscopic analyses of cells collected from a *B. subtilis* 3610 pellicle biofilm bearing dual fluorescent reporters of P*_comGA_-gfp* and P*t_apA_-mKate2* (EH43). The activity of P*_tapA_-mKate2* (cells in red) indicates expression of the key biofilm matrix operon *tapA-sipW-tasA* while P*_comGA_-gfp* reports a late competence gene *comGA* (cells in green). More images are available in Supple. Fig 2. Scale bar, 10 μm. Scale bar is representative to all images in the figure. **(B)** Flow cytometry analyses of the above dual fluorescent reporter strain (EH43, indicated as mKate2/GFP), two single reporter strains (QS34 for P*_comGA_-gfp* and EH41 for P*_tapA_-mKate2*, indicated as GFP and mKate2, respectively), and 3610 (as a gating control). Activities of P*_comGA_-gfp* and P*_tapA_-mKate2* were measured in the GFP (y-axis) and RFP (for mKate2, x-axis) filters, respectively. Numbers represent the percentage (%) of gated cells vs total cells in the corresponding quadrant. **(C)** The quadrant analyses of the flow cytometry results. The percentage indicates gated cells/total cells in corresponding quadrant. Each dot represents one biological replicate. Experiments were repeated three times. Error bars indicate standard deviations. [One dot representing the mKate2/GFP quadrant from the single reporter (EH41, shown as GFP) and one dot representing the mKate2/GFP quadrant from the double reporter (EH43, shown as mKate2/GFP) were omitted due to errors].

### Overexpression of the competence activator gene *comK* blocks biofilm formation in *B. subtilis*

We hypothesized that the biofilm and the competence pathways may negatively cross-regulate each other and consequently these two cell types become mutually exclusive. To test our hypothesis, we first looked at the competence pathway. Since ComA-ComP is also important for biofilm formation (49), we focused on the competence activator ComK, which acts downstream of ComA-ComP in the pathway (Fig. 1)(44). To test if ComK negatively regulates the biofilm pathway, a Δ*comK* deletion mutant was constructed and the biofilm phenotype of the mutant examined. Surprisingly, the mutant did not show any noticeable biofilm phenotype compared to the wild type (Fig. 5A). Given the ultralow ratio of cells expressing P*_comGA_-gfp* (Fig. 4A) and the knowledge on ComK regulation, we reasoned that ComK was not active in the majority of cells, which could explain why Δ*comK* and the wild type strain did not differ in the biofilm phenotype. We then tested *comK* overexpression. An IPTG-inducible copy of *comK* was constructed and introduced into 3610. This time, upon addition of IPTG, the engineered strain displayed a strong biofilm defect (Fig. 5A), similar to what was seen in the laboratory strain 168 (Fig. 2A). This indicates that ComK strongly impacts biofilm development in *B. subtilis*. ComK activation is known to eventually cause growth arrest in *B. subtilis* (50). As an important control, the growth of the *comK* overexpression strain was examined. Upon induction of *comK* in the presence 10 μM IPTG, no difference in the growth rate of the cells was found when compared to without *comK* induction (Supple. Fig. 3A).

**Figure 5.**
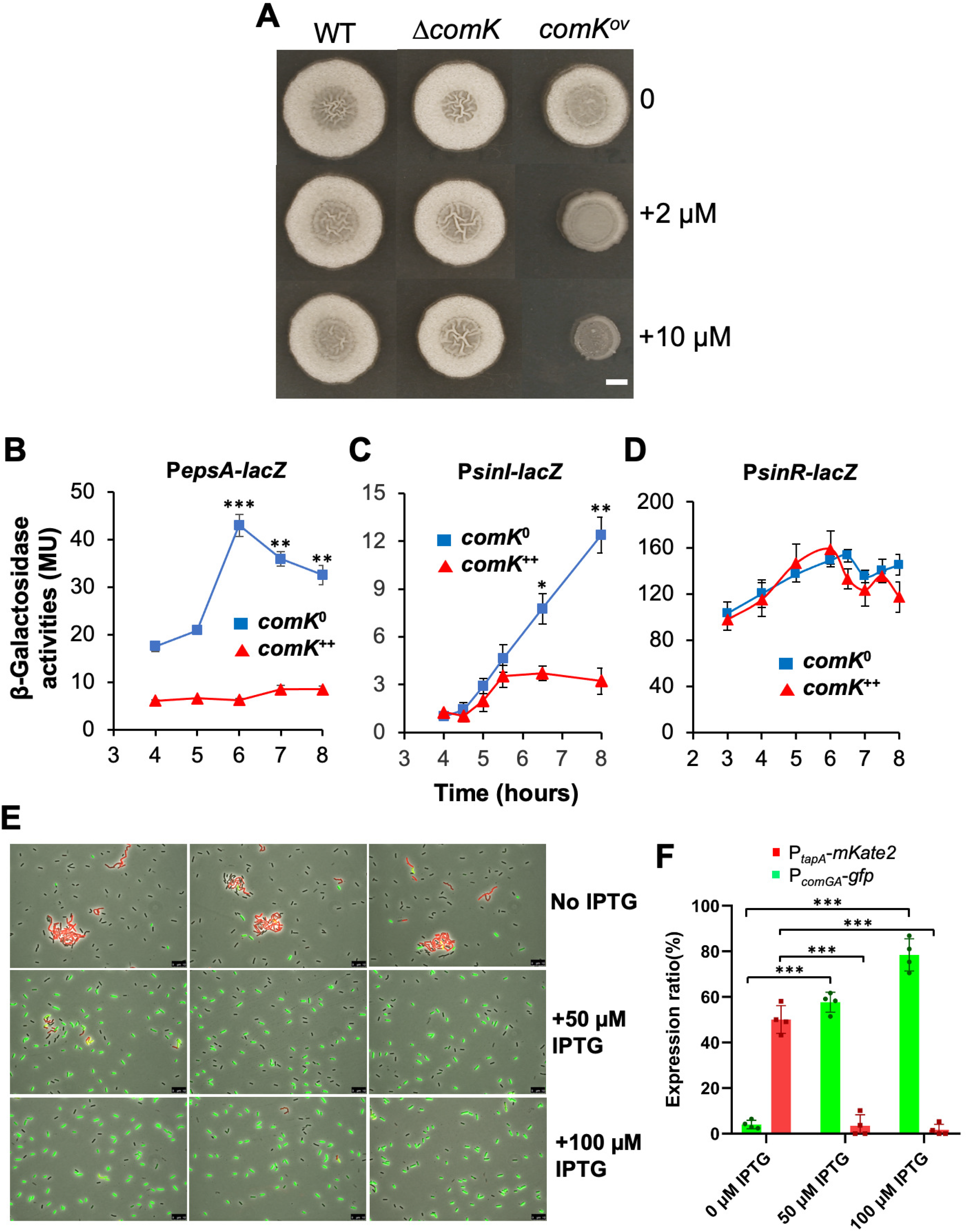
*comK* negatively regulates key biofilm genes. **(A)** Overexpression of *comK* impairs biofilm formation in *B. subtilis*. Phenotypes of the colony biofilms by the wild type (3610), Δ*comK* (YC100), and the wild type strain harboring an IPTG-inducible copy of *comK* (YC142) on MSgg plates supplemented with 0, 2, or 10 μM IPTG. Scale bar, 2 mm. The scale bar is representative for all pictures in this panel. **(B-D)** Wild type strains bearing both an IPTG inducible copy of *comK* and one of the three biofilm gene reporters, P*_epsA_-lacZ* **(B,** YC160), P*sM-lacZ* **(C,** YC159) and P*_sinR_-lacZ* **(D,** YC177), were assayed for ß-galactosidase activities. Cells were cultured in MSgg in shaking, in the absence (*comK*^0^) or presence (*comK*^++^) of 10 μM IPTG to induce *comK*. Assays were done in triplicates. Error bars represent standard deviations. The *t*-test was applied for statistical analysis. * indicates P value < 0.05, ** indicates P value < 0.005, and *** indicates P value < 0.0005. **(E)** Fluorescent microscopic analyses of the dual reporter strain (P*_comGA_-gfp* and P*_tapA_-mKate2*) that also contains an IPTG inducible copy of *comK* (EH44). Cells were grown in shaking MSgg to log phase (OD_600_=0.5), split into three fractions, one without IPTG and the other two with either 50 or 100 μM IPTG, and continued to grow for an hour before harvested and analyzed under fluorescent microscopy. Scale bars, 10 μm. **(F)** Quantitative analyses of the dual reporter activities upon *comK* overexpression. For each IPTG concentration (0, 50, or 100 μM), individual dots represent results from 4 separate images (in one biological replicate) comprising of about 600-800 cells in total. Error bars represent standard deviations. The *t*-test was applied for statistical analysis. *** indicates P value < 0.0005.

### ComK negatively regulates biofilm matrix genes

To further characterize the impact of ComK on biofilm formation, we tested if ComK regulates any of the matrix genes, such as the *epsA-O* and the *tapA* operons. A previously constructed transcription reporter P*_epsA_-lacZ* was introduced into the *comK* overexpression strain (51). The resulting strain (YC160) was used to test the impact of *comK* overexpression on the activity of P*_epsA_-lacZ*. As shown in Fig. 5B, the activity of P*_epsA_-lacZ* decreased dramatically upon addition of 10 μM IPTG to induce *comK*. In another test, we introduced the *comK* overexpression construct (*thrC*::P*_spank_-comK*) into the previously described dual fluorescent reporter strain (P*_tapA_-mkate2* and P*_comGA_-gfp*). When the resulting strain (EH44) was grown in MSgg without addition of IPTG, about 50% of the cells were expressing P*_tapA_-mkate2* (cells in red, Fig. 5E-F), and again a very low number of cells expressed P*_comGA_-gfp* (cells in green, Fig. 5E-F). Upon addition of 50 or 100 μM IPTG to induce *comK* expression for about an hour, the competence reporter P*_comGA_-gfp* was found activated in a significantly increased subpopulation of cells (this time about 58% or 78% of the total cells were in green, middle and lower panels, respectively, Fig. 5E-F) while the expression of P*_tapA_-mKate2* was turned off almost completely (cells in red, middle and lower panels, Fig. 5E-F). Importantly, matrix producers and competent cells again rarely overlapped in this assay. These results suggest that ComK negatively regulates biofilm matrix genes.

### ComK negatively regulates *sinI*

The *epsA-O* and the *tapA* operons are directly repressed by SinR while derepression occurs when SinI counteracts SinR through protein-protein interactions (23, 24, 52, 53). To determine how ComK negatively regulates the matrix operons, we tested if ComK regulates either *sinI* or *sinR*. We took a similar approach by applying previously constructed transcription reporters of P*_sinI_-lacZ* and P*_sinR_-lacZ* (51). Each reporter was introduced into the *comK* overexpression strain, and the impact of *comK* overexpression on the activity of the reporters was similarly tested. Indeed, *comK* overexpression was found to have a strong negative impact on the activity of P_*sinI*_-*lacZ*, but not on P*_sinR_-lacZ* (Fig. 5C-D). These results indicate that ComK negatively regulates the matrix operons likely through its regulation on *sinI*.

### ComK directly binds to the regulatory region of *sinI*

ComK regulates genes through binding to the so-called K-box often found in the regulatory region of the genes (54). When the promoter sequence of *sinI* was analyzed, a region of DNA sequence that resembles the consensus K-box (“AAAA-N_5_-TTTT-N_8_-AAAA-N_5_-TTTT”) was recognized (Fig. 6A). This DNA sequence overlaps with both the −35 and −10 motives of the sigma A-dependent promoter and a Spo0A~P activation site (OA~P) in the *sinI* promoter (33). ComK binding to this putative K-box could prevent *sinI* transcription.

**Figure 6.**
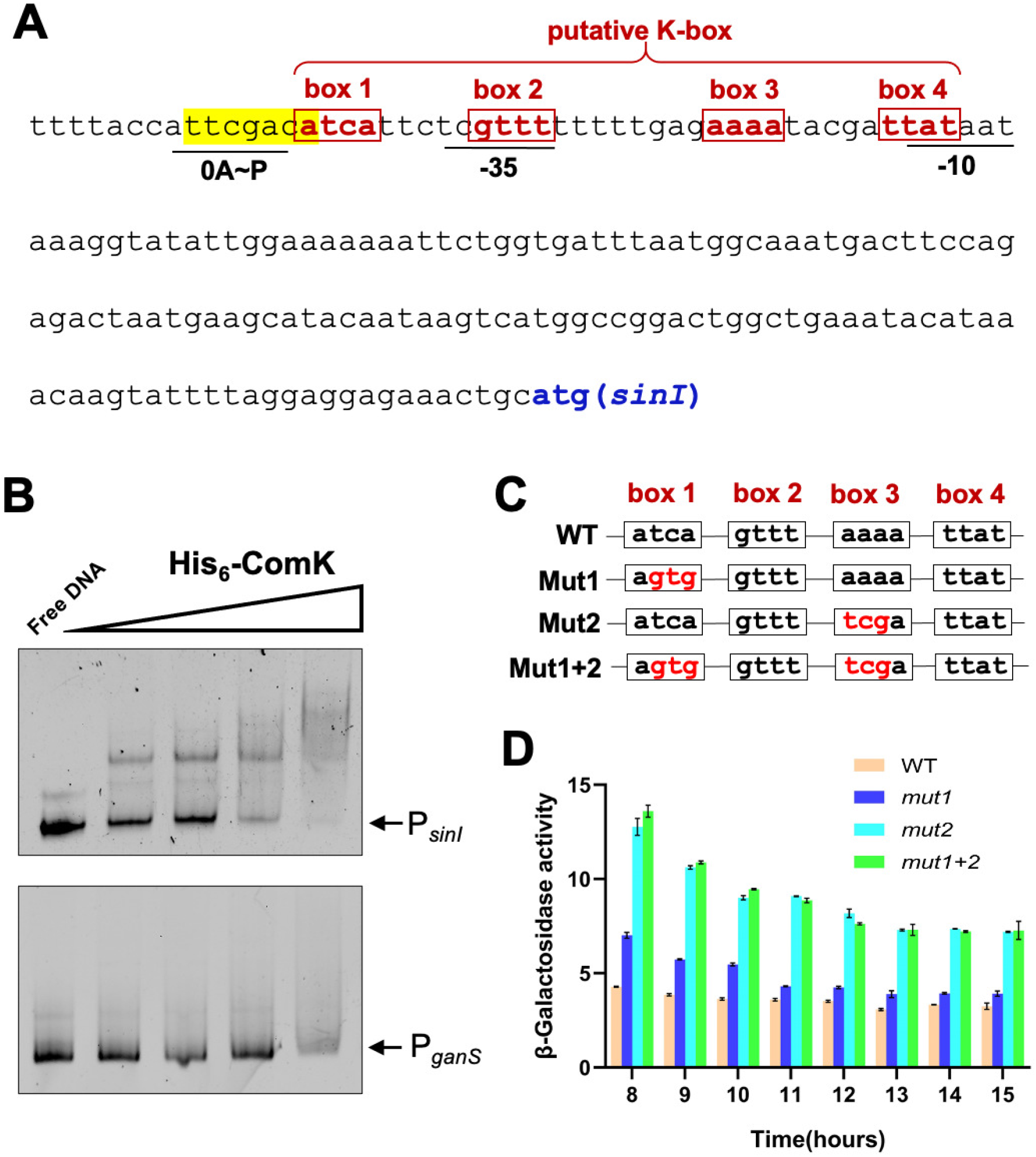
ComK directly binds to the promotor of *sinI*. **(A)** Shown is the DNA sequence of the promoter region of *sinI*. The Spo0A~P activation site (in yellow and underlined), the −10 and −35 motives (underlined), and putative ComK boxes (from box 1 to 4) are highlighted (31). **(B)** Electrophoretic mobility shift assay (EMSA) of His_6_-ComK binding to the promoter of *sinI. P_sinI_* was end-labeled with Cy3 dye and used as the DNA probe. Cy3-labeled P_*ganS*_ was used as a negative control. The far-left lanes are the control of free DNA without proteins. A decreasing gradient of 150, 60, 15, and 7.5 nM of the recombinant His_6_-ComK was applied in the lanes in EMSA as indicated. 200 pmol of fluorescent labelled DNA probe was applied in each lane. **(C)** Site-directed mutagenesis of the ComK boxes in the *sinI* promoter is indicated. Nucleotide changes in box 1 (mut1) and box 3 (mut2) are highlighted in red. Changes in box 2 and box 4 are avoided due to their overlap with −10 and −35 promoter motives. **(D)** β-Galactosidase activities of the cells with an inducible *comK* construct and bearing either wild type P*_sinI_-lacZ* or the reporter fusions with indicated point mutations in the K-box (mut1, mut2, and mut1+2, as shown in **C**) were performed. IPTG was added at 10 μM in the media. Cells were grown in shaking MSgg. Samples were periodically collected and assayed for β-galactosidase activities. Assays were performed at least in triplicates. Error bars represent standard deviations.

To test if the ComK protein binds to the promoter of *sinI*. An electronic mobility shift assay (EMSA) was performed. Recombinant His_6_-ComK proteins were expressed in *E. coli* and purified. Fluorescently end-labeled DNA probe containing about 300 bp of the *sinI* promoter was mixed with a gradient of His_6_-ComK proteins in the mobility shift assay. ComK was found to shift the DNA fragment, indicating a direct binding (upper panel, Fig. 6B). As a negative control, a similar size DNA probe containing the promoter of a *ganS* gene, not known to be regulated by ComK (54, 55), was used in the same assay and little DNA shift was observed (lower panel, Fig. 6B). Thus, the binding of ComK to the *sinI* promoter appeared to be specific. To further test if ComK recognizes the putative K-box in the *sinI* promoter, site-directed mutagenesis was performed on the box1 and box3 of the putative K-box as indicated (Fig. 6C, nucleotide changes highlighted in red). Mutagenesis on box2 and box4 was avoided due to their overlap with the −35 and −10 promoter motives (Fig. 6A). The reporter strains bearing P*_sinI_-lacZ* with sited directed mutations in the K-box were constructed and the activities of those strains tested. The results show that the point mutations in the box3 (mut2) and in both boxes 1 and 3 (mut1+2) had the most significant effect, resulting in increased *sinI* expression (Fig. 6D). To summarize, the competence pathway negatively cross-regulates the biofilm pathway likely through the competence activator ComK directly repressing the key biofilm regulatory gene *sinI*.

### Biofilm matrix negatively impacts competence in *B. subtilis*

We also predicted that the biofilm pathway negatively regulates competence development. A previous study showed that the extracellular biofilm matrix physically blocked cells from sensing the competence pheromones, which is essential for ComA-ComP-mediated activation of the *srfAA-AD* operon (49). The *srfAA-AD* operon harbors the key competence gene *comS*, critical for activating ComK and inducing late competence genes in *B. subtilis* (42). Here, we further showed that deleting the biofilm matrix genes (Δ*epsH*Δ*tasA*) indeed improved competence of *B. subtilis* 3610; the transformation efficiency of the matrix double mutant was ~7 fold higher than that of the wild type (Fig. 7A). This phenomenon was not only seen in 3610, but also in some other environmental isolates of *B. subtilis*. When the *epsA-O* operon was deleted in those environmental isolates and the transformation efficiency of the mutants was compared to the respective wild type strains, an increase in transformation efficiency from about 2 to 100 folds was seen in 6 out of the 9 strains (Fig. 7B and Supple. Fig. 3B). This result indicates that it may be a general mechanism that the presence of extracellular matrix can reduce the competence of *B. subtilis* cells.

**Figure 7.**
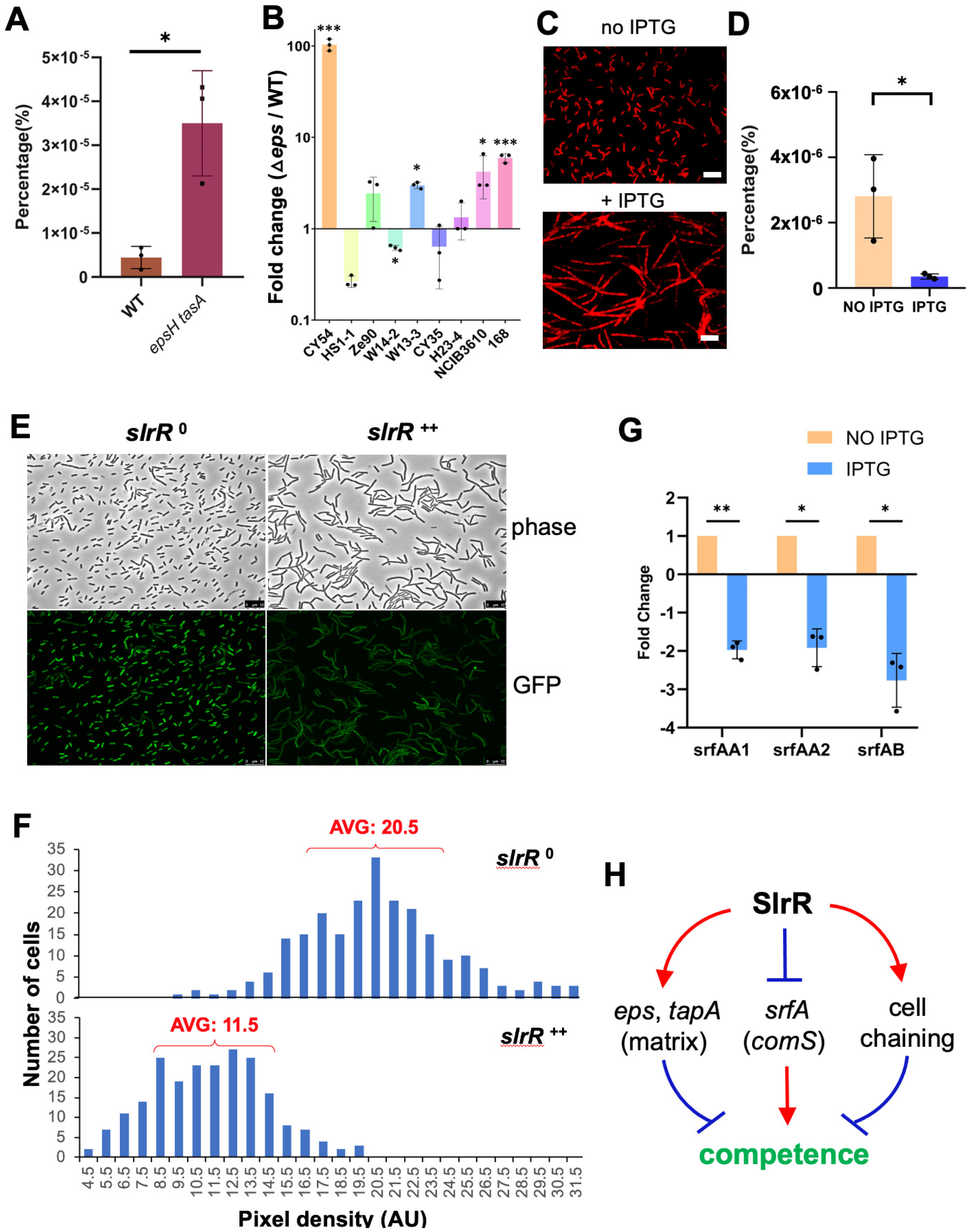
SlrR negatively regulates competence through three distinct mechanisms. **(A)** Comparison of the transformation efficiency of the wild type (3610) and the Δ*epsHΔtasA* double mutant (YC775). Results were presented as percentage (%) of the number of transformants relative to the total number of cells. Experiment was repeated three times. Each dot indicates one biological replicate. Error bars indicate standard deviation. * indicates P value < 0.05. **(B)** Comparison of the transformation efficiency between the 7 environmental strains of *B. subtilis* and their respective Δ*epsA-O* mutants (4). Strains 168, 3610 and the Δ*epsA-O* mutants of 168 and 3610 were also included. Results were shown as fold changes of the CFU counts during transformation comparing the Δ*epsA-O* mutants and the respective wild type strains. Each dot indicates one biological replicate. Error bars indicate standard deviation. * indicates P value < 0.05 and *** indicates P value < 0.0005. **(C)** The microscopic images of a *slrR* inducible strain (YC672). Cells were grown in shaking LB without or with the addition of 100 μM IPTG to induce *slrR* expression and cell chaining phenotype. Red indicates cell membrane staining by the membrane dye FM 4-64. Scale bars, 10 μm. **(D)** Comparison of the transformation efficiency of the *slrR* inducible strain (YC672) in the absence or presence of 100 μM IPTG. The transformation efficiency is shown as percentage of the number of transformants vs the total number of cells. The experiment was repeated three times. Each dot represents one biological replicate. Error bars indicate standard deviation. * indicates P value < 0.05. **(E)** Fluorescence microscopic analyses of the *slrR* overexpression strain harboring a fluorescent reporter of P*_srfAA_-gfp* (YC1270) in the absence or presence of IPTG to induce *slrR* expression. Cells were grown in shaking MSgg to early log phase (OD_600_=0.3), and split into two fractions, one without IPTG (*slrR*^0^) and the other with 100 μM IPTG (*slrR^++^*) added to induce *slrR* expression for an hour before harvest and analysis of the cells. **(F)** Quantification of fluorescent pixel density of the cells in **(E)** by ImageJ (with MicroJ plugin). More than 200 cells from each sample were randomly picked for analysis. The results were plotted indicating the difference in P*_srfAA_-gfp* activity between without and with *slrR* overexpression. The numbers 20.5 (*slrR*^0^) and 11.5 (*slrR^++^*) indicate average pixel density (AU) of the top 50% of the cells in each population. **(G)** qPCR analyses to test the negative regulation of SlrR on *srfAA-AD*. Total RNA was prepared from the *slrR* inducible strain (YC672) grown with (*slrR^++^*) and without (*slrR*^0^)100 μM IPTG. Three primer pairs, two for detection of *srfAA* and one for detection of *srfAB*, were applied. Each experiment was repeated three times. Each dot indicates one biological replicate. The error bars indicate standard deviation. * indicates P values < 0.05. ** indicates P value < 0.005. **(H)** A schematic drawing of how the biofilm regulator SlrR negatively impacts competence through three distinct mechanisms. In Figure 7, the *t*-test was applied for statistical analysis.

### Extensive chaining in the biofilm negatively impacts competence

Cells in the *B. subtilis* biofilm form long bundled chains, important for building organized 3-dimentional structure of the biofilm (Fig. 4A)(19, 56). Interestingly, the DNA uptake machinery was shown to localize to the poles of the cells during *B. subtilis* competence development (57, 58). If true, one would predict that the nonmotile chained cells may encounter reduced efficiency in DNA uptake. To test the possible impact of chaining on competence, we first applied a Δ*sigD* mutant. Sigma D (SigD) is responsible for the transcription of genes encoding multiple autolysins for cell separation, but not known to directly influence competence (59). The Δ*sigD* mutant formed extensive long chains, consistent with the previous report (Supple. Fig. 4A)(59). Interestingly, when the transformation efficiency was compared between the wild type and the Δ*sigD* mutant, the mutant showed drastically reduced efficiency even after taking into consideration of the impact of cell chains on CFU counting (see methods). The diminished competence in Δ*sigD* was almost comparable to that of the Δ*comK* mutant (Supple. Fig. 4B).

In the wild type biofilm, chaining is also controlled by the SinR/SlrR switch (26, 30, 32, 56, 59). SlrR^ON^ cells tend to form long chains of cells bundled together by the extracellular matrix due to matrix genes being activated while autolysin genes being simultaneously shut off (26). We decided to test if increasing SlrR production by gene overexpression could have a similar negative impact on the transformation efficiency seen in Δ*sigD*. A previously constructed Δ*slrR* mutant containing an IPTG inducible copy of *slrR* was applied (56). Both the chaining phenotype and the transformation efficiency of the engineered cells were compared between no addition and with addition of IPTG. As shown in Figs. 8C-D, adding IPTG to the media significantly increased cell chaining even in shaking conditions while substantially reduced the transformation efficiency compared to no addition of IPTG. Our results thus suggest that extensive cell chaining during *B. subtilis* biofilm development likely plays a role in limiting competence of *B. subtilis* cells.

### SlrR negatively regulates the *srfAA-AD* operon

A previous study showed that the null mutation of *sinR* abolished competence but not clear how (60). Since the inhibition on competence by Δ*sinR* was reported in a laboratory strain unable to form robust biofilms, overproduction of the matrix by the Δ*sinR* cells may not be the answer. Here we present evidence that SlrR, whose gene is repressed by SinR, negatively regulates *srfAA-AD* and thus competence (Fig. 1). We introduced a P*_sfAA_-gfp* fluorescent reporter into the *slrR* mutant bearing an IPTG inducible copy of *slrR*. Upon addition of IPTG, *slrR* was induced, evidenced by the chaining phenotype and the activity of P*_srfAA_-gfp* noticeably decreased compared to no addition of IPTG (Fig. 8E). Fluorescent pixel density in individual cells was quantified using ImageJ. In cells overexpressing *slrR*, the average pixel density was about half of that in cells not overexpressing *slrR* (11.5 vs 20.5, Fig. 8F). The repression of *srfAA-AD* by SlrR was also confirmed by real-time quantitative PCR using three different probes for the operon (Fig. 8G). Our genome-wide transcription profiling to characterize global SlrR regulon showed similar results (Chai, unpublished). Our results thus suggest that SlrR negatively regulates the *srfAA-AD* operon. In summary, we believe that SlrR, together with SinR, negatively regulates competence through three distinct mechanisms (Fig. 8H), by i) promoting matrix production to block competence signaling, and ii) forming extensive cell chains to possibly block DNA uptake, and iii) negatively regulating the *srfAA-AD* operon (and conceivably *comS*, Fig. 1).

## Discussion

Environmental strains of *B. subtilis* have important applications in agriculture, industry, and biotechnology. However, different from well-studied laboratory strains, those environmental strains are genetically less accessible for still unclear reasons. This imposes limitation of their application in various fields. Environmental strains of *B. subtilis* are often capable of forming robust biofilms both under the laboratory settings and on plant roots, an ability important for them to establish intimate relationship with the plant in the rhizosphere. In this study, we uncovered that during biofilm formation by the *B. subtilis* model strain 3610, a very low number of cells differentiates into competent cells. Similar observations were also made in several environmental strains of *B. subtilis* (Fig. 3). We presented evidence that the low competence is contributed by the ability to form robust biofilms. The very low ratio of competent cells in the *B. subtilis* biofilm might be ecologically more relevant than what is often studied in test tubes with optimized competence media. The biological implication of greatly reduced competence associated with biofilm formation by wild *B. subtilis* strains is not very clear. One could argue that this is how natural competence in *B. subtilis* is expected to function in order to balance the ability of generating genetic variations and the potential risk of having too many individual cells in the population acquiring genetic variations.

Cell differentiation is a hall mark feature of bacterial biofilm formation. A number of studies have demonstrated co-existence of distinct cell types in the *B. subtilis* biofilm, and in a few cases have characterized molecular mechanisms of how specific cell types become mutually exclusive or inclusive (2, 14, 26, 28, 29). In this study, we showed that competent cells and matrix producers are mutually exclusive cells types during *B. subtilis* biofilm formation and provided mechanistic explanations of why they become mutually exclusive. Based on our evidence, we propose a working model on the cross-regulation between these two developmental pathways that contributes to their mutual exclusivity (Fig. 1). In one regulation, ComK in the K-state cells directly turns off expression of the key biofilm gene *sinI*. Since activity of *sinI* is indispensable for biofilm activation, direct repression of *sinI* by ComK allows K-state cells to shut down the biofilm pathway, and eliminate matrix production and cell chaining, which we showed negatively influence transformation efficiency. In the other regulation, the biofilm regulator SlrR plays a central role that allows SlrR^ON^ cells to avoid competence development. Not only matrix may physically block competence quorum-sensing, but cell chaining and negative regulation of the *srfAA-AD* operon (and *comS*) by SlrR further contribute to the shut-off of competence in SlrR^ON^ cells. Similar regulations could also be present in the *B. subtilis* environmental strains. However, since we carried out our mechanistic studies primarily in the model strain 3610, further studies will be needed to test those regulations directly in the environmental strains of *B. subtilis*. In addition, the regulation in those environmental strains could differ from that in 3610, as some of our data already indicate so (e.g. Fig. 7B).

Previous studies have shown that competent cells are inclusive to cells responding to the competence pheromones and activating the surfactin biosynthesis operon, which very intriguingly also transcribes a small *comS* gene key to the competence development (42, 49). According to a previous study, under biofilm conditions in 3610, the genes involved in initiating competence (e.g. *comQ-comX-comP*) is expressed in a majority of the cells, while the surfactin operon is expressed in only about 10% of the total number of cells (defined as *s*f^N^) in the population (49). This significant difference in the ratio is thought to be contributed in part by the paracrine signaling mechanism in that although the majority of cells produce the competence pheromone, only a handful of them respond to this quorum-sensing signal. Further, another peptide pheromone CSF is also involved by regulating the response regulator ComA through a feedback mechanism (61). It can be speculated that in those 10% *s*f^N^ cells, ComK is activated above a critical threshold in a further reduced ratio of the cells, due to the well-studied complex regulation on ComK and the *comK* gene. Those ComK^ON^ cells enter the so-called K-state and ultimately become competent for environmental DNA acquisition. It is a bit surprising to us that the ComK^ON^ cells (presumably cells expressing P*_comGA_-gfp*) is less than 0.1%, versus 10% of *s*f^N^ cells. This implies that at best only about 1 out of 100 cells enter the K-state even after all the cells initiate the competence by inducing the *srfAA-AD* operon and *comS*. Again, it could be evolutionarily important to limit competence capacity when *B. subtilis* cells live in multicellular communities in the natural environment.

## Materials and Methods

### Strains and media

Strains used in this study are listed in Table 1. *B. subtilis* strains PY79, 168, NCIB3610, their derivatives, and environmental isolates of *B. subtilis* were cultured in lysogeny broth at 37°C. Pellicle biofilm formation in *B. subtilis* was induced using MSgg broth (50 mM potassium phosphate and 100 mM MOPS at pH 7.0 supplemented with 2 mM MgCl_2_, 700 μM CaCl_2_, 50 μM MnCl_2_, 50 μM FeCl_3_, 1 μM ZnCl_2_, 2 μM thiamine, 0.5% glycerol, and 0.5% glutamate) at 30°C. Colony biofilm formation of *B. subtilis* was induced using MSgg solidified with 1.5% (w/v) agar at 30°C. When growing the chromosomal *thrC* integration strains of *B. subtilis*, additional 300 μg ml^-1^ threonine was added. Enzymes used in this study were purchased from New England Biolabs (MA, USA). Chemicals and reagents were purchased from Sigma or Fisher Scientific (MA, USA). Oligonucleotides were purchased from Eurofins Genomics (PA, USA) and DNA sequencing was also performed at Eurofins Genomics. Antibiotics, if needed, were applied at the following concentrations: 5 μg ml^-1^ of tetracycline, 1 μg ml^-1^ of erythromycin, 100 μg ml^-1^ of spectinomycin, 10 μg ml^-1^ of kanamycin, and 5 μg ml^-1^ of chloramphenicol for transformation in *B. subtilis* and 100 μg ml^-1^ of ampicillin and 50 μg ml^-1^ of kanamycin for *E. coli* DH5α and BL21/DE3 strains.

**Table 1.** Strains used in this study.

| Strain | Genotype | Reference |
| --- | --- | --- |
| ***B. subtilis*** |  |  |
| PY79 | SPβ-cured laboratory strain of *B. subtilis* as a host for transformation | (68) |
| 168 | A laboratory strain of *B. subtilis* as a host for transformation |  |
| NCIB3610 | Undomesticated *B. subtilis* strain capable of biofilm formation | (19) |
| EH41 | *sacA::P<sub>tapA</sub>-mKate2* in 3610, kan<sup>R</sup> | This study |
| EH43 | A dual fluorescent reporter strain of *sacA::P<sub>tapA</sub>-mKate2* and *amyE::P<sub>comGA</sub>-gfp* in 3610, km<sup>R</sup>, cm<sup>R</sup> | This study |
| EH44 | *sacA::P<sub>tapA</sub>-mKate2*, *amyE::P<sub>comGA</sub>-gfp*, *thrC::P<sub>hpspank</sub>-comK* in 3610, km<sup>R</sup>, cm<sup>R</sup>, erm<sup>R</sup> | This study |
| RL4169 | *ΔsigD::tet* in 3610, tet<sup>R</sup> | (59) |
| YC100 | *ΔcomK::kan* in 3610, km<sup>R</sup> | This study |
| YC108 | *amyE::P<sub>sinR</sub>-lacZ* in 3610, cm<sup>R</sup> | (51) |
| YC110 | *amyE::P<sub>sinI(WT)</sub>-lacZ* in 3610, cm<sup>R</sup> | (51) |
| YC130 | *amyE::P<sub>epsA</sub>-lacZ* in 3610, cm<sup>R</sup> | (51) |
| YC157 | *amyE::P<sub>hpspank</sub>-comK* in 3610, spec<sup>R</sup> | This study |
| YC159 | *thrC::P<sub>hpspank</sub>-comK* and *amyE::P<sub>sinI</sub>-lacZ* in 3610, erm<sup>R</sup>, cm<sup>R</sup> | This study |
| YC160 | *thrC::P<sub>hpspank</sub>-comK* and *amyE::P<sub>epsA</sub>-lacZ* in 3610, erm<sup>R</sup>, cm<sup>R</sup> | This study |
| YC177 | *thrC::P<sub>hpspank</sub>-comK* and *amyE::P<sub>sinR</sub>-lacZ* in 3610, erm<sup>R</sup>, cm<sup>R</sup> | This study |
| YC672 | *ΔslrR::tet*, *amyE::P<sub>hpspank</sub>-slrR*, in 3610, tet<sup>R</sup>, spec<sup>R</sup> | (26) |
| YC775 | *ΔepsH::tet* and *ΔtasA::spec* in 3610, tet<sup>R</sup>, spec<sup>R</sup> | This study |
| YC1270 | *amyE::P<sub>hpspank</sub>-slrR*, *lacA::P<sub>srfAA</sub>-gfp* in 3610, spec<sup>R</sup>, erm<sup>R</sup> | This study |
| QS34 | *amyE::P<sub>comGA</sub>-gfp* in 3610, cm<sup>R</sup> | This study |
| QS35 | *thrC::P<sub>spank</sub>-comK*, *amyE::P<sub>sinI(WT)</sub>-lacZ* in 3610, mls<sup>R</sup>, cm<sup>R</sup> | This study |
| QS36 | *thrC::P<sub>spank</sub>-comK*, *amyE::P<sub>sinI(mut1)</sub>-lacZ* in 3610, mls<sup>R</sup>, cm<sup>R</sup> | This study |
| QS37 | *thrC::P<sub>spank</sub>-comK*, *amyE::P<sub>sinI(mut2)</sub>-lacZ* in 3610, mls<sup>R</sup>, cm<sup>R</sup> | This study |
| QS38 | *thrC::P<sub>spank</sub>-comK*, *amyE::P<sub>sinI(mut1+2)</sub>-lacZ* in 3610, mls<sup>R</sup>, cm<sup>R</sup> | This study |
| QS42 | *ΔdegQ::spec* in Ze90, spec<sup>R</sup> | This study |
| QS43 | *ΔdegQ::spec* in HS1-1, spec<sup>R</sup> | This study |
| QS44 | *ΔdegQ::spec* in CY35, spec<sup>R</sup> | This study |
| QS45 | *ΔdegQ::spec* in CY54, spec<sup>R</sup> | This study |
| QS46 | *ΔdegQ::spec* in W13-3, spec<sup>R</sup> | This study |
| QS47 | *ΔdegQ::spec* in W14-2, spec<sup>R</sup> | This study |
| QS48 | *ΔdegQ::spec* in H23-4, spec<sup>R</sup> | This study |
| CY35 | An environmental isolate of *B. subtilis* | (4) |
| CY54 | An environmental isolate of *B. subtilis* | (4) |
| H23-4 | An environmental isolate of *B. subtilis* | (4) |
| HS1-1 | An environmental isolate of *B. subtilis* | (4) |
| W13-3 | An environmental isolate of *B. subtilis* | (4) |
| W14-2 | An environmental isolate of *B. subtilis* | (4) |
| Ze90 | An environmental isolate of *B. subtilis* | (4) |
| CY94 | *ΔdegU* in 3610, tet<sup>R</sup> | This study |
| CY96 | *ΔdegS* in 3610, tet<sup>R</sup> | This study |
| CY601 | *ΔdegQ::spec* in 3610, spec<sup>R</sup> | This study |
| CY602 | *degQ* overexpression in 3610, spec<sup>R</sup> | This study |
| ***E. coli*** |  |  |
| DH5α | An *E. coli* host for molecular cloning | Invitrogen |
| QS16 | *E. coli* BL21/DE3 with the plasmid pQS06 | This study |
| **Plasmids** |  |  |
| pYC119 | *amyE::P<sub>hpspank</sub>-comK* in pDR111, amp<sup>R</sup>, spec<sup>R</sup> | This study |
| pYC121 | *amyE::gfp* (promoterless) in pDG1662, amp<sup>R</sup>, sm<sup>R</sup> | (51) |
| pYC128 | *thrC::P<sub>hpspank</sub>-comK* in pDG1664, amp<sup>R</sup>, erm<sup>R</sup> | This study |
| pYC166 | *amyE::P<sub>sinI</sub>-lacZ* in pDG268, amp<sup>R</sup>, cm<sup>R</sup> | (33) |
| pQS06 | pET28a(P<sub>T7</sub>-*his6-comK*) plasmid, kan<sup>R</sup> | This study |

### Strain construction and DNA manipulation

General methods for molecular cloning followed the published protocols (62). Restriction enzymes (New England Biolabs) were used according to the manufacturer’s instructions. Transformation of plasmid DNA into *B. subtilis* strains was performed as described previously (63). SPP1 phage-mediated general transduction was also used to transfer antibiotic-marked DNA fragments among different strains (64). Plasmids used in this study are listed in Table 1 and oligonucleotides are listed in Table 2.

**Table 2.**
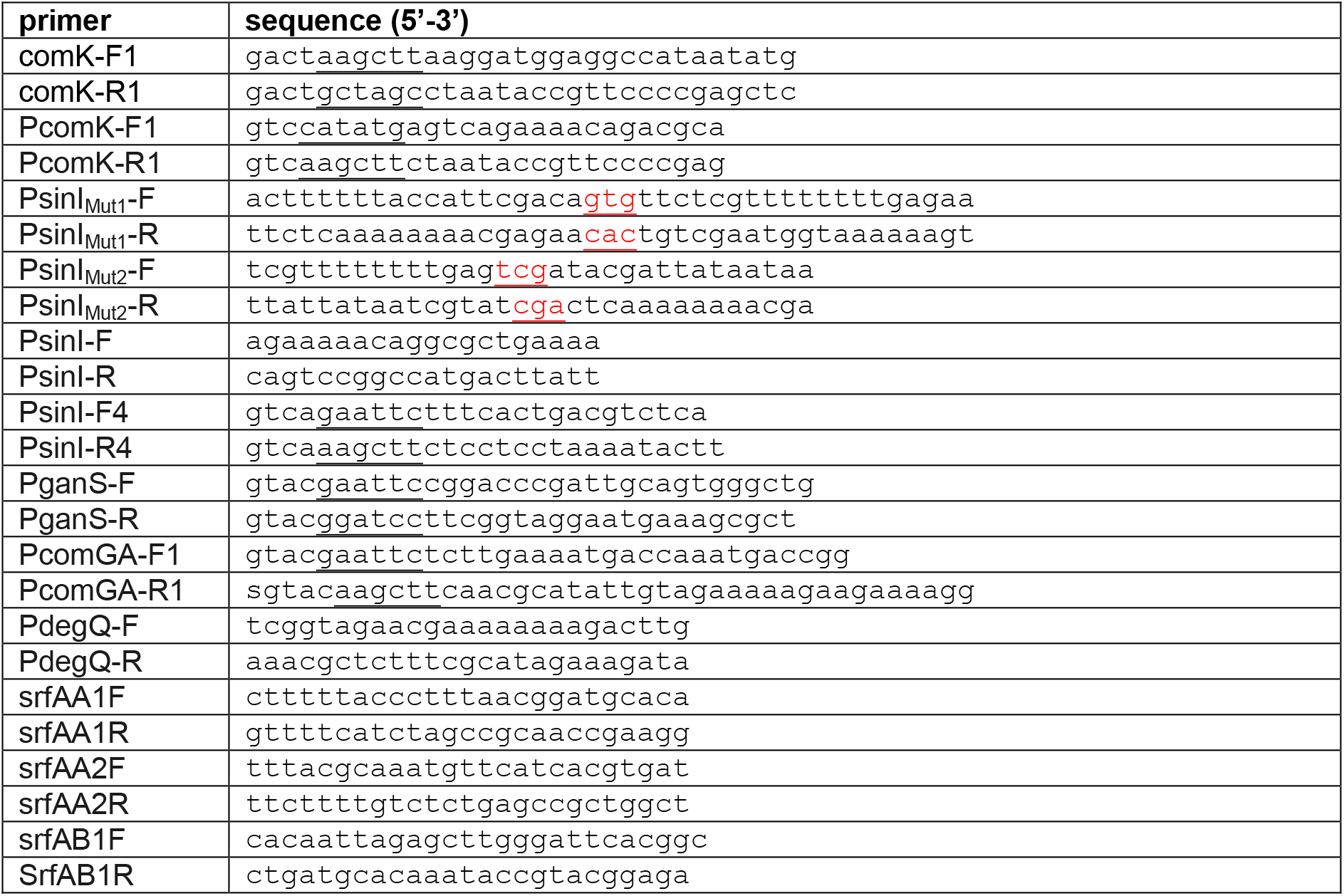
Oligonucleotides used in this study.

To generate the competence gene reporter strains (P*_comGA_-gfp*), the promoter of *comGA* was amplified via PCR using 3610 genomic DNA as the template and primers PcomGA-F1 and PcomGA-R1. The PCR product was cloned into the EcoRI and HindIII sites of pYC121 aiming for the integration into the *amyE* locus of 3610 and other environmental isolates of *B. subtilis*. The recombinant plasmid was transformed into DH5α for amplification. The recombinant plasmid extracted from transformed DH5α was subsequently transformed into PY79 and then to 3610.

To generate *comK* insertional deletion mutation in 3610 (YC100), the lysate containing Δ*comK::kan* was made from RL2262 (a gift from Rich Losick, Harvard University) and introduced into 3610 by transduction. To create an IPTG-inducible copy of *comK* for integration at the *amyE* locus, the *comK* coding sequence was amplified by PCR using primers comK-F1 (HindIII) and comK-R1 (NheI). The PCR product was digested and cloned into the HindIII and NheI sites of pDR111 (65) to make an IPTG inducible P*_spank_-comK* fusion, generating a recombinant plasmid pYC119. The pYC119 plasmid was then used for integration of P*_spank_-comK* into the *amyE* locus of 3610. To do so, the plasmid was first introduced into PY79 by transformation and then into 3610 by SPP1 phage mediated transduction. To create a second version of an IPTG-inducible *comK* for integration at the *thrC* locus of 3610, a DNA fragment containing the P*_spank_* promoter was cut from the above pYC119 with EcoRI and HindIII double digestion, and a second DNA fragment containing the *comK* coding sequence and the *lacI* gene was cut separately from pYC119 by HindIII and BamHI double digestion. These two DNA fragments were cloned into the EcoRI and BamHI sites of pDG1664 by three-way ligation to generate an IPTG-inducible P*_spank_-comK* in the *thrC* integration plasmid, resulting in pYC128. The pYC128 plasmid was introduced into PY79 by transformation. To generate three reporter strains of YC159 (P*_sinI_-lacZ*), YC160 (P*_epsA_-lacZ*), and YC177 (P*_sinR_-lacZ*), each with an inducible copy of *comK*at the *thrC* locus, the lysate containing *thrC::P_spank_-comK::mls* was prepared from the above recombinant pY79 strain, and introduced into YC108 (P*sinR-lacZ*), YC110 (P*_S_sinI-lacZ*), and YC130 (P*epsA-lacZ*), respectively, by SPP1 phage mediated transduction.

To generate the recombinant plasmid pQS06 for His_6_-ComK overexpression and purification, the *comK* coding gene was amplified by PCR using 3610 genomic DNA as the template and primers PcomK-F1 and PcomK-R1. The PCR product was cloned into the pET28a vector between the restriction sites NdeI and HindIII to create the PT7-his6-*comK* fusion. The recombinant plasmid pQS06 was prepared from *E. coli* DH5α and then introduced into *E. coli* BL21/DE3 by chemical transformation. The resulting *E. coli* strain QS16 was used for His6-ComK overexpression and purification. To create the strain YC1270, the lysate containing *amyE*::P*_hyperspank_-slrR* was prepared from YC672 and introduced into DL744, which bears the reporter *lacA::P_srfAA_-gfp::mls*, by transduction (49, 56). To create the transcription reporter fusion of P*_comGA_-gfp*, the promoter sequence of the *comGA* gene was amplified by PCR using 3610 genomic DNA as the template and primers PcomGA-F1 and PcomGA-R1. The PCR product was cloned into the pYC121 plasmid between the restriction sites EcoRI and HindIII to create the P*_comGA_-gfp* fusion. The recombinant plasmid was transformed into DH5α for amplification. The recombinant plasmid extracted from transformed DH5α was subsequently transformed into PY79 and then to 3610.

### Colony and pellicle biofilm development

For colony biofilm formation, cells were grown to exponential phase in LB broth and 2 μL of the culture was spotted onto MSgg media solidified with 1.5% (w/v) agar. The plates were incubated at 30°C for 3 days. For pellicle biofilm formation, cells were grown to exponential phase in LB broth, and 3 μL of the culture was inoculated into 3 mL of MSgg liquid media in a 6-well or 12-well microtiter plate (VWR). The plates were incubated at 30°C for 2-3 days. Images of colony and pellicle biofilms were taken using a Nikon Coolpix camera or a Leica MSV269 dissecting scope.

### Site-directed mutagenesis

Site-directed mutagenesis of the *sinI* regulatory sequence was performed by using 6 different primers to change the nucleotides in the putative ComK binding boxes in the *sinI* promoter region. In the P_*sinI*_^Mut1^ construction (box 1), PsinI-F4 and PsinI^Mut1^-R were used to amplify the fragment 1. PsinI-R4 and PsinI^Mut1^-F were used for the amplification of fragment 2. The fragments 1 and 2 were subsequently used in the second round of overlapping PCR to generate the full-length DNA fragment containing the *sinI* promoter with designated point mutations. Overlapping PCR product was purified using PCR purification kit (Qiagen) and subsequently digested using EcoRI and HindIII. The plasmid pDG268 was digested simultaneously using EcoRI and HindIII. Both digestion products were gel-purified, ligated using T4 Ligase, and transformed *E. coli* DH5α. The recombinant plasmid was purified from *E. coli* DH5α and transformed into PY79. The *amyE* homologous region containing mutated P_*sinI*_^Mut1^ and the chloramphenicol resistance marker was integrated onto PY79 chromosome via double crossover homologous recombination. The genomic DNA of the resulting transformant was prepared and subsequently transformed into 3610. Site-directed mutagenesis on P_*sinI*_^Mut2^ (box3) was performed similarly except that the primers PsinIMut2-F, PsinIMut2-R, PsinI-F4 and PsinI-R4 were used. Construction of P_*sinI*_^Mut1+2^ was performed by using the recombinant plasmid containing P_*sinI*_^Mut1^ as the template during the first round of PCR amplification and primers PsinIMut2-F, PsinIMut2-R, PsinI-F4 and PsinI-R4. Designated point mutations in the *sinI* promoter on the recombinant plasmids were verified by DNA sequencing before being introduced into *B. subtilis*.

### Assays of transformation efficiency

Assays on transformation efficiency were performed by introducing *B. subtilis* genomic DNAs containing specific antibiotic resistance genes as a selection marker into indicated strains. Specifically, the three plasmids pDG1662 (*amyE*::*chl*^R^), pDG1663 (*thrC*::*mls*^R^), and pDG1730 (*amyE*::*spec*) containing different antibiotic markers flanked by the either *B. subtilis amyE* or *thrC* sequences (26), were introduced into 3610 first for double crossover recombination on the chromosome. The genomic DNA bearing either *amyE*::*chl*^R^, or *amyE*::*chl*^R^, or *thrC*::*mls* was prepared from the above strains. The concentration of the prepared genomic DNAs was determined using Nanodrop (Thermo Fisher). For each transformation event, a fresh single colony of the strain was picked and grown in LB broth to log phase. The log phase culture was then 1:100 subcultured into 2 mL competence medium (MC) supplemented with 3 mM MgSO_4_. Cells were grown at 37 °C in shaking until early stationary phase (OD_600_=1.5), 10 μg of the genomic DNA was then mixed with 500 μL competent cells, and cells were incubated for another hour before harvest. Samples were plated on the LB plates with the addition of either 100 μg/mL of spectinomycin (for *spec* selection) or 5 μg/mL of chloramphenicol (for *chl*^R^ selection) or 25 μg/mL of lincomycin+1 μg/mL of erythromycin (for *mls*^R^ selection). Next day, CFU on the transformation plates was counted. The total number of cells was calculated by measuring the O.D._600_ of the culture prior to plating and assuming 3 x 10^8^ cells for OD_600_=1.0 of the culture (which was experimentally determined for 3610, data not shown) across all *B. subtilis* cultures used in the transformation assays unless for the strains involving extensive cell chains (see below). Each assay was done at least three times.

For transformation of the *slrR* inducible strain (YC672), cells were grown in LB broth to log phase, diluted 1:100 into MSgg broth, and grown to mid log phase (OD_600_= 0.5) again. Cells were then split into two fractions, one added with 100 μM IPTG to induce *slrR* overexpression and the other no addition of IPTG. Both cultures were continued to grow for another hour for *slrR* induction. 10 μg of genomic DNA was then added to each culture of 500 μL followed by one more hour growth at 37°C with shaking. Before harvest, the cultures were mildly sonicated (scale 1.5 output, 50% interval, 3-5 pulses on ice) using the sonicator (Scienz). After sonication, cells were analyzed under light microbiology to verify the disruption of chaining. Cells were then the plated on LB plates supplemented with appropriate antibiotics. Next day, the number of transformants on the plates were calculated. For counting the total number of cells, cultures were serial-diluted and plated on regular LB plates. CFU was counted next day. All assays were done at least three times with biological replicates. The transformation experiment using the *sigD* mutant followed the similar protocol to eliminate the impact of chaining.

### Expression and purification of recombinant ComK proteins

BL21/DE3 cells harboring the recombinant plasmid (PT7-his6-*comK*) were grown in LB broth supplemented with 50 μg/mL kanamycin at 37°C overnight in shaking. The overnight culture was aliquoted at 1:500 to 300 mL LB media supplemented with 50 μg/mL kanamycin in shaking condition at 30 °C. 1 mM IPTG was added when OD_600_ of culture reached 0.5. IPTG induction was allowed to continue for two hours before harvesting the culture. The culture was harvested and centrifuged at 4500 rpm at 4 °C for 30 minutes. The cell pellet was resuspended and washed twice using cold phosphate buffer solution. The supernatant was discarded, and the cell pellets were again resuspended using 10 mL lysis buffer (20 mM Tris-HCl, 300 mM NaCl, 1 mM PMSF, pH 8.5). The cell resuspension was lysed using sonication on ice. The total cell lysate was centrifuged at 5000 rpm at 4 °C for 30 minutes. The cleared lysate containing soluble His6-ComK was transferred into a new precooled tube. Cell lysate was mixed with 1 mL of Ni-NTA agarose beads (Qiagen), and the mixture was rotated at 4 °C for 2 hours. The mixture of lysate and beads was transferred into the column and washed for five times with wash buffer (20 mM Tris-HCl, 300 mM NaCl, 25 mM imidazole, pH 8.5). 2 mL wash buffer was applied in each wash. The flow-through was also collected in 5 separate tubes. The column was eluded five times using elution buffer (20 mM Tris-HCl, 300 mM NaCl, 250 mM imidazole, pH 8.5). 500 μL of elution buffer was applied to the column each time and the elute was collected in the tubes separately. 12% SDS-PAGE was applied to size-fractionate the proteins and verify the purity and abundance of the recombinant His6-ComK proteins. The purified protein fractions were pooled and dialyzed in a dialysis buffer (20 mM sodium phosphate, 300 mM NaCl, 0.3 mM DTT, 10% glycerol, pH 7.4) overnight. The final concentration of the protein was determined using Bradford protein assays. The proteins were stored at −80 °C.

### Electrophoretic mobility shift assays (EMSA)

An about 300-bp DNA fragment containing the promoter of *sinI* (P_*sinI*_) was used as the DNA probe for the binding of recombinant ComK proteins in EMSA, and a similar size DNA fragment containing the promoter of *ganS* (P_*ganS*_) was used as a negative control DNA probe. The fluorescent DNA probes were generated by PCR using 3610 genomic DNA as the template, and with the forward primers P_*sinI*_-F and P_*ganS*_-F covalently linked to 5’Cy3 fluorescent dye, and the regular reverse primers P_*sinI*_-R and P_*ganS*_-R. The PCR product was gel purified, eluded in ddH_2_O and the quality was measured by NanoDrop (Fisher Thermo Scientific). A gradient of protein concentrations was applied in the reaction mixtures. A decreasing gradient of 150, 60, 15, and 7.5 nM of the recombinant His_6_-ComK was applied in each binding mixture. 200 pmol of fluorescent labelled DNA probe was applied in each lane. The protein-DNA binding reaction was incubated in 20 μL reaction volume, containing 10 mM Tris-HCl, 10 mM HEPES, 50 mM KCl,1 mM EDTA, 10 μg/mL BSA, and 4% sucrose. To reduce non-specific binding, 500 ng of random DNA (poly dI:dC) was added to each binding reaction. The reaction was incubated on ice for 30 min. The gel was run in 0.5X TBE buffer at 65 V for 3.5 h at 4 °C. The resulting gel was imaged using ChemiDoc MP (Bio-Rad, USA).

### Assays of β-galactosidase activity

Cells were cultured in MSgg medium at 30°C with shaking. When indicated, IPTG was added to the media at the beginning at a final concentration of 10 μM. One milliliter of culture was collected at each indicated time point and cells were centrifuged down at 5,000 rpm for 10 min. Cell pellets were suspended in 1 ml Z buffer (40 mM NaH_2_PO_4_, 60 mM Na_2_HPO_4_, 1 mM MgSO_4_, 10 mM KCl, and 38 mM β-mercaptoethanol) supplemented with 200 μg ml^-1^ lysozyme. Resuspensions were incubated at 37°C for 15 min. Reactions were started by adding 200 μl of 4 mg ml^-1^ ONPG (2-nitrophenyl-β-D-galactopyranoside) and stopped by adding 500 μL of 1 M Na_2_CO_3_. Samples were then briefly centrifuged down at 5,000 rpm for 1 min. The soluble fractions were transferred to cuvettes (VWR), and absorbance of the samples at 420 nm was recorded using a Bio-Rad Spectrophotometer. The β-galactosidase specific activity was calculated according to the equation (Abs_420_/time×OD_600_) × dilution factor × 1000. Assays were conducted in triplicate.

### Cell membrane staining

For cell membrane staining, cells were grown to log phase and harvested. Cell pellets were washed with PBS buffer twice, resuspended in 100 μL PBS buffer, and mixed with 1 μL of FM 4-64 dye (Life Technologies) for 5 min on ice with gentle tapping of the tube. 2 μL of the resuspension was placed on a 1% (w/v) agarose pad and covered with a cover slip. For observation of the FM 4-64 fluorescence dye, the the excitation wavelength was set at 540-580 nm and the emission wavelength at 610-680 nm. Cells from three independent biological replicates were imaged using a Leica DFC3000 G camera on a Leica AF6000 microscope.

### Real time quantitative PCR (qPCR)

Cells were collected after the overexpression of SlrR in experimental group. Total RNAs were extracted by using TRIzol (Invitrogen) following the manufacturer’s protocol. Isolated RNAs were reverse transcribed into single-stranded complementary DNA (cDNA) using a High Capacity cDNA Reverse Transcription Kit (Applied Biosystems). RT-qPCR was performed by using Fast SYBR™ Green Master Mix (Applied Biosystems) with Step One Plus Real-Time PCR system (Applied Biosystems). The 16S rRNA gene was used as an internal reference. The relative expression of specific genes was calculated by using the 2^-ΔΔCT^ method. The statistically analysis was performed using t-test.

### Cell fluorescence imaging and pixel quantification

For imaging of the environmental strains bearing the P*_comGA_-gfp* fluorescent reporter, the reporter strains were grown in LB broth to log phase. Each log phase culture was then 1:100 subcultured into 2 mL competence medium (MC) supplemented with 3 mM MgSO_4_. Cells were grown at 37 °C in shaking until early stationary phase (OD_600_=1.5). Cells were then spun down, washed with PBS buffer once, and resuspended in 100 μL of PBS buffer. 2 μL of the resuspension was placed on a 1% (w/v) agarose pad, covered with a cover slip, and observed under fluorescent microscopy. Imaging of different samples was conducted using the same exposure settings.

To quantify the ratio of P*_comGA_-gfp* expressing cells relative to the total number of cells, fluorescence of single cells was quantified in 3 different images comprising of a total of 600-800 cells per sample, using the MicrobeJ plugin for ImageJ (66, 67). These 3 images were randomly selected from more than half-dozen separate images obtained in two experimental repeats. Using the “Analyze Particles” command, the size cutoff of 100 pixel^2 was set to exclude the noise from the viable cells. Using the “Threshold” command, the threshold number was adjusted to highlight and select the pixel area of interest, which indicates the viable cells, for the analysis. A threshold above three times of the average background pixel density was used to define P*_comGA_-gfp* expressing cells. The total number of cells were counted in phase images while the fluorescent cells were counted with the corresponding fluorescent channel images and verified in combination with manual examination.

For imaging of the *slrR* inducible strain (YC1270), cells were grown in LB broth to log phase. Cells were then diluted 1:100 to MSgg broth and grown to mid log phase (OD_600_= 0.5). Cells were then split into two fractions, one added with 100 μM IPTG and the other no addition of IPTG. Both fractions were continued to grow for another hour. Cells were then spun down, washed with PBS buffer once, and resuspended in 100 μL of PBS buffer. 2 μl of the resuspension was placed on a 1% (w/v) agarose pad, covered with a cover slip, and observed under fluorescent microscopy. Non-specific background fluorescence was determined by quantifying WT cells bearing no fluorescent reporter. Imaging of different samples was conducted using the same exposure settings. Pixel density of single-cell fluorescence was quantified on >200 cells per sample using the MicrobeJ plugin for ImageJ.

### Flow cytometry

Flow cytometry was carried out using a BD FACSAria II with a 70 micron nozzle. Briefly, biofilms were grown for 48 hours in defined monosodium glutamate-glycerol (MSgg) biofilm promoting-media. After 48 hours of growth, cells were harvested from pellicle biofilms, cell chains were disrupted by mild sonication, and 5 μL of resuspended cells were diluted in 1 mL PBS through a 35 μm filter (Corning Falcon tube, Thermo Fisher). FACS DIVA software was used to collect 100,000 events for each sample. Data were analyzed in FlowJo software. Gates were drawn, based on the size, to exclude the bulky events, which were considered as clumps or cell chains. This size bias was confirmed by plotting these events on FSC/SSC axis. The rest of the gated cells was considered to be single cells and were displayed in GFP-A/mKate-A axis. Four strains, 3610 as a gating control for fluorescent signals, two single reporter strains, (P*_comGA_-gfp* and P*_tapA_-mkate2*), and the dual reporter strain (P*_comGA_-gfp/ Pt_apA_-mkate2*), were applied in the analyses by flow cytometry. Assays were performed in three biological replicates.

## Acknowledgments

This work was supported by the US National Science Foundation (MCB1651732 to Y. Chai), and by the National Natural Science Foundation of China (31922074), National Key Research and Development Program of China (2017YFD0201104), and the Young Elite Scientist Sponsorship Program (2017QNRC001) to Y. Chen.

## Supplemental Figure Legends

**Supple. Figure 1. DegQ impacts competence negatively in some, but not all, environmental strains. (A)** Transformation efficiency of WT, Δ*degQ, degQ* overexpression, Δ*degS*, and Δ*degU* mutants of *B. subtilis* 3610. Results were shown as percentage of the number of transformants vs the number of total cells. Assays were done in triplicates. Error bars represent standard deviations. * indicates P value < 0.05. The *t*-test was applied for statistical analysis. **(B)** Comparison of transformation efficiency between the Δ*degQ* deletion mutants and the respective wild type environmental strains. Numbers in the y-axis represent fold changes comparing CFU counts of the Δ*degQ* deletion mutants and those of the respective wild type strains. Assays were done in triplicates. Error bars represent standard deviations. **(C)** DNA sequence alignments of the *degQ* promoter region from 7 environmental strains plus 168 and 3610. Newly identified nucleotide changes in environmental strains, which correspond to the −28, −48, and −77 positions relative to the *degQ* transcription start, are highlighted in red boxes. The previously identified single nucleotide change in 168 is highlighted in the blue box.

**Supple. Figure 2. Matrix producers and competent cells are mutually exclusive in the 3610 biofilm.** Fluorescent microscopic analyses of cells collected from a *B. subtilis* 3610 pellicle biofilm bearing dual fluorescent reporters of P*_comGA_-gfp* and P*t_apA_-mKate2* (EH43). In the above overlay images, the activity of P*_tapA_-mKate2* (cells in red) indicates expression of the key biofilm matrix operon *tapA-sipW-tasA* while P*_comGA_-gfp* reports a late competence gene *comGA* (cells in green). Scale bar, 10 μm. Scale bar is representative to all images in the figure.

**Supplemental Figure 3. (A) Mild overexpression of *comK* does not cause growth inhibition.** The wild type strain harboring an IPTG-inducible copy of *comK* (YC142) was inoculated in MSgg broth and grown at 37°C over a period of 10 hours. IPTG was either not added (*comK*^0^) or added to the medium at the final concentration of 10 μM to induce mild *comK* overexpression (*comK*^++^). Cell were periodically collected and cell optical density (O.D._600_) was measured. (B) Comparison of transformation efficiency between the *epsA-O* mutants of the environmental strains and their respective wild type strains. Results were presented in log scale as percentage of the number of transformants vs the number of total cells. Experiment was done in triplicates. Error bars represent standard deviations. * indicates P value < 0.05; *** indicates P value < 0.0005. The *t*-test was applied for statistical analysis.

**Supplemental Figure 4. Δ*sigD* cells form long chains and demonstrate very low transformation efficiency. (A)** The microscopic images of the wild type (3610) cells and the Δ*sigD* mutant (RL4169), known to form long cell chains due to lack of SigD-controlled autolysin activities. Red indicates cell membrane staining by the membrane dye FM 4-64. Scale bar, 10 μm. Scale bar applies to both images here. **(B)** Comparison of transformation efficiency of the wild type (3610) and the Δ*sigD* mutant (RL4169). The Δ*comK* mutant (YC100) is known to be deficient in transformation and used as a negative control in the experiment. Assays were done in triplites. Error bars represent standard deviations. The *t*-test was applied for statistical analysis. ** indicates P value < 0.005.

